# A Reduced Mechanistic Model for Aquatic Decomposition of a Human Body

**DOI:** 10.64898/2026.09.29.755382

**Authors:** Ufuk Akın, Ömer Faruk Aykır, Neslihan Nesliye Pelen, Melike Kaplan

**Author notes:** & (Neslihan Nesliye Pelen).

## Abstract

Estimating the postmortem submersion interval (PMSI) remains challenging due to complex biological and environmental interactions during aquatic decomposition. This study proposes a reduced mechanistic ordinary differential equation (ODE) model linking tissue degradation, microbial activity, and dissolved oxygen dynamics. Calibrated against the total aquatic decomposition score (TADS) trajectory for Northern Adriatic Sea cases, the model preserves a strictly monotone TADS–time relationship, ensuring unique numerical inversion from observed TADS to PMSI.

Synthetic observation experiments evaluated how data selection influences parameter recovery, practical identifiability, and robustness under measurement uncertainty. Microbial measurements improved parameter recovery for microbial dynamics, whereas dissolved oxygen measurements constrained oxygen-consumption and renewal processes. Combining both types of observation optimized the recovery of coupled parameters. Trajectory reconstruction, profile likelihood analysis, boundary-hit frequencies, and Monte Carlo simulations confirmed these findings.Importantly, while all observation strategies reproduced TADS with comparable accuracy, they differed substantially in recovering underlying biological parameters and latent-state dynamics.

Thus, accurately reproducing decomposition scores alone is insufficient for reliable parameter inference. This framework provides a transparent and biologically interpretable foundation for the mechanistic estimation of PMSI. Future applications to case-level and multi-site datasets may enable site-specific recalibration and improved predictive evaluation across diverse aquatic environments.

## 1 Introduction

Universal formulations describing terrestrial decomposition processes have been proposed in the literature [1]. However, when postmortem decomposition occurs in aquatic environments, the temporal progression and morphological characteristics observed in terrestrial settings undergo substantial modification [2]. Aquatic contexts introduce additional sources of variability, including fluctuating immersion conditions, delayed recovery, postmortem drift, and scene-specific constraints, all of which contribute to increased uncertainty in estimating the postmortem submersion interval (PMSI) [3, 4, 5]. These complexities underscore the need for structured, environment-specific approaches in forensic practice.

In routine casework, detailed environmental measurements – such as continuous dissolved oxygen profiles, comprehensive microbial analyses, or complete environmental time series –are rarely available at the time of body recovery. Consequently, modeling frameworks capable of operating with incomplete yet routinely available forensic information are particularly relevant.

The rate and pattern of aquatic decomposition are influenced by salinity, microbial activity, hydrodynamics, scavenger activity, and climatic conditions [4]. While terrestrial decomposition stages have been extensively characterized, aquatic-specific modifications such as adipocere formation and maceration limit the direct applicability of land-based models to submerged bodies [1]. Several aquatic decomposition scoring systems have therefore been proposed to provide standardized descriptions of decomposition progression under submerged conditions, including the systems introduced by Heaton et al., van Daalen et al., and subsequent validation studies in both marine and freshwater environments [3, 6, 7, 5]. To address documentation challenges in such contexts, semi-quantitative scoring systems—including FAD, BAD, LAD, and TADS—have been developed to provide reproducible and scene-independent characterization of decomposition status, thereby facilitating communication among forensic pathologists, investigators, and courts [3, 6, 4, 7, 5]. The case series presented by Palazzo et al. [4] demonstrated the applicability of TADS under specific environmental conditions.

Although empirical score–time regressions derived from such case series are useful within the contexts in which they were established, similar empirical approaches based on accumulated degree-days (ADD) and decomposition scoring have also been widely investigated for terrestrial and aquatic postmortem interval estimation [8, 9, 10]. More recently, probabilistic modeling approaches have also been proposed to integrate decomposition observations with environmental variables for postmortem interval estimation [11, 12]. Likewise, mathematical models have been successfully applied to other areas of forensic science, including postmortem body cooling and heat-transfer analysis [13]. However, mechanistic dynamical models explicitly describing the biological progression of aquatic decomposition and linking these processes to decomposition score evolution have received comparatively little attention. As a result, the direct transferability of empirical models to distinct aquatic environments may be limited. In this setting, a process-oriented modeling framework may offer a complementary perspective by linking observed score dynamics to underlying biophysical influences in an abstract and mathematically structured manner. The proposed framework represents decomposition at the level of macroscopic score dynamics rather than detailed microbiological or biochemical mechanisms.

Figure 1 summarizes the conceptual structure of the proposed framework. Environmental forcing enters the mechanistic ODE system through temperature-dependent tissue degradation and oxygen limitation. The latent tissue-loss variables are then mapped to regional aquatic decomposition scores, which are summed to obtain TADS. Finally, monotonicity of the calibrated TADS trajectory enables inverse estimation of PMSI from an observed decomposition score.

**Figure 1.**
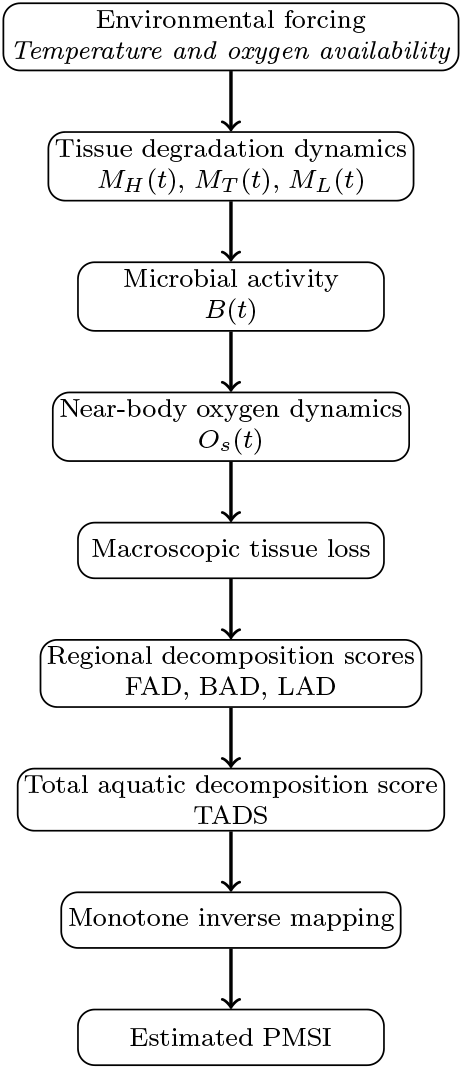
Conceptual workflow of the proposed reduced mechanistic framework. Environmental forcing modulates tissue degradation, microbial activity, and near-body oxygen dynamics. The resulting tissue-loss trajectory is mapped to regional decomposition scores and then to TADS. When the calibrated TADS– time relationship is monotone, numerical inversion provides an estimate of the postmortem submersion interval (PMSI).

The proposed framework supports PMSI assessment by providing a transparent and interpretable mapping between observed decomposition scores and elapsed submersion time. By establishing a strictly monotone score–time relationship, the model guarantees a unique numerical inversion from TADS to PMSI, thereby eliminating potential ambiguities in forensic reporting. Beyond introducing a mechanistic ODE framework, this study systematically investigates how different biologically relevant observation strategies influence parameter recovery, practical identifiability, and calibration robustness. Together, these contributions establish an extensible mathematical foundation for PMSI estimation that can be systematically calibrated across diverse aquatic environments and provide guidance for future mechanistic studies using richer biological and environmental observations.

## 2 Methods

The overall methodology comprises environmental module configuration, calibration against the published TADS–PMSI relationship, monotonicity assessment, and the subsequent inverse estimation of PMSI. The governing differential equations for tissue dynamics, microbial activity, and oxygen depletion utilized throughout this framework are formally formulated in Section 3.

### 2.1 Calibration overview

The present study provides a proof-of-concept calibration of the reduced mechanistic ODE model against the published aggregated regression between total aquatic decomposition score (TADS) and postmortem submersion interval (PMSI) reported by Palazzo et al. [4]. Publicly available aquatic decomposition datasets are currently limited to aggregated score–time relationships, whereas individual case-level observations are generally unavailable or incomplete. Consequently, the published TADS–PMSI regression provides the most comprehensive publicly available quantitative reference for calibrating the proposed mechanistic model. The empirical regression function

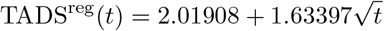

(with *t* in days) is treated as a macroscopic calibration target.

Rather than fitting to individual case-level PMSI observations (which are not publicly available), the model is calibrated so that its predicted score trajectory TADS^model^(*t*) approximates TADS^reg^(*t*) over a specified time interval.

### 2.2 Fitting interval and time discretization

Calibration is performed over the interval

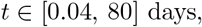

consistent with the range illustrated in [4].

A logarithmically spaced time grid

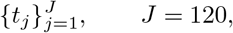

is used in order to balance early-time and late-time behavior of the trajectory.

### 2.3 Environmental forcings and module configuration

In accordance with the Northern Adriatic Sea case descriptions, scavenger activity is assumed negligible and the scavenger module is deactivated:

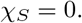

The oxygen module is activated:

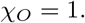

Water temperature *T*_env_(*t*) is prescribed as an external environmental forcing consistent with the case descriptions in [4].

Bulk dissolved oxygen concentration *O*_∞_(*t*) is prescribed as a constant representative value within the reported Northern Adriatic Sea range (approximately 6–9 mg L^−1^), based on independent regional oceanographic studies (e.g. [14]). The near-body oxygen concentration *O*_*s*_(*t*) is determined endogenously by the oxygen box model.

### 2.4 Numerical integration

The ODE system is integrated forward in time from *t* = 0 using an adaptive Runge–Kutta method of order 4(5) (RK45). Absolute and relative tolerances are chosen sufficiently small to ensure numerical stability across the fitted time interval.

All tissue fractions *M*_*i*_(*t*) are required to remain nonnegative throughout integration. Parameter combinations leading to nonphysical (negative or unstable) trajectories over the fitting interval are rejected during optimization.

### 2.5 Observation model

Regional scores are defined via a shared linear mapping. Since the Total Aquatic Decomposition Score (TADS) is defined as the sum of the three regional decomposition scores (FAD, BAD, and LAD), each having a minimum possible score

of 1, the observation-model intercept is fixed at this minimum value. In the absence of region-specific calibration data, a common slope parameter is assumed for all anatomical regions to obtain a parsimonious parameterisation.

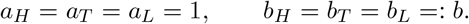

Thus,

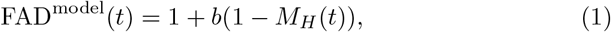

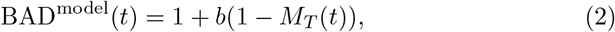

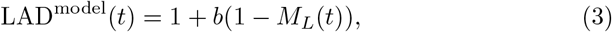

and

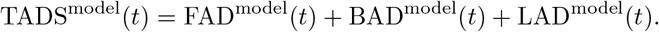

To avoid over-parameterisation, and because region-specific calibration data are unavailable for the aggregated regression target, only a common slope parameter is estimated across anatomical regions.

### 2.6 Objective function and parameter estimation

The reduced parameter vector to be estimated is

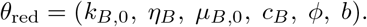

Parameters are estimated by minimizing the unweighted sum of squared errors

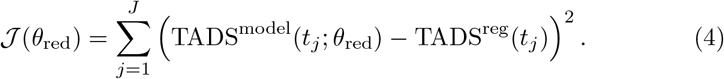

Given the aggregated nature of the calibration target, the optimization seeks an effective parameterization reproducing the observed score trajectory rather than a unique biological identification of every model parameter. Optimization is performed under positivity constraints for all rate parameters. Multiple randomized initializations are used in order to reduce sensitivity to local minima, and the parameter set achieving the smallest objective value is retained.

### 2.7 Monotonicity and inverse mapping

After calibration, the fitted trajectory TADS^model^(*t*) is verified to be monotonically increasing over [0.04, 80]. When monotonicity holds, inversion is performed numerically by solving

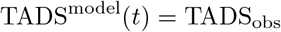

for *t* within the calibrated interval.

The inverse mapping is therefore defined only within the fitted range and under the assumed environmental forcings.

### 2.8 Scope and limitations of the calibration

The present calibration is tied to the Northern Adriatic Sea context represented by [4]. Estimated parameters should therefore be interpreted as effective, environment-specific quantities rather than universal biological constants. Application to other aquatic environments would require appropriate environmental characterization and re-calibration.

## 3 Model formulation

This section describes the mathematical structure of the proposed reduced dynamical system developed to simulate the macroscopic progression of aquatic decomposition. To capture the main biophysical drivers while avoiding excessive parameterisation, the framework integrates tissue degradation, microbial activity, and near-body oxygen dynamics within a coupled ordinary differential equation (ODE) system. Furthermore, the model adopts a modular architecture that allows selected biological components to be activated or deactivated depending on the environmental characteristics of the forensic context.

### 3.1 Reduced ODE model with optional scavenger and oxygen modules

In order to avoid over-parameterisation while still retaining the main mechanistic drivers of aquatic decomposition, we construct a reduced dynamical model in which scavenger effects and oxygen limitation can be optionally activated depending on the environmental context.

We introduce two binary switches:

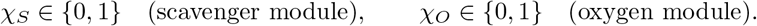

When scavenging activity is negligible (as reported for the Northern Adriatic Sea cases), we set *χ*_*S*_ = 0. In environments where scavenger activity is expected to play a significant role, *χ*_*S*_ = 1. Similarly, when dissolved oxygen data are unavailable or oxygen limitation is assumed negligible, we set *χ*_*O*_ = 0; otherwise *χ*_*O*_ = 1.

The state variables of the reduced model are

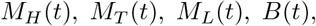

and, when *χ*_*O*_ = 1, the near-body oxygen concentration *O*_*s*_(*t*).

#### 3.1.1 Model parameters and their interpretation

The parameters appearing in the model equations have the following biological and physical interpretations (Table 1).

**Table 1:** Model parameters for the reduced mechanistic aquatic decomposition model. All parameters listed below appear explicitly in the governing equations. Parameters are grouped into fixed quantities (set from general biochemical principles or structural considerations), environmental forcings prescribed independently, and effective parameters estimated from decomposition score data.

| Parameter | Role in the model | Value / treatment | Justification and references |
| --- | --- | --- | --- |
| **(i) Fixed parameters (not estimated)** |  |  |  |
| $T_{\text{ref}}$ | Reference temperature for $Q_{10}$ scaling | Fixed (e.g. 10°C or 20°C) | Standard reference in temperature-dependent rate models |
| $Q_{10,i}$<br>($i = H, T, L$) | Temperature sensitivity of tissue degradation rates | Fixed: $Q_{10,i} = 2$ | Empirical temperature dependence of enzymatic and microbial processes [15, 16, 17] |
| $Q_{10,B}$ | Temperature sensitivity of microbial activity | Fixed: $Q_{10,B} = 2$ | Microbial metabolic rates approximately double per 10°C [15, 16, 17] |
| $Q_{10,\mu}$ | Temperature sensitivity of microbial loss rate | Fixed: $Q_{10,\mu} = 2$ | Consistent thermal scaling of biochemical reaction rates [15, 16] |
| $M_{i,0}$ | Initial tissue fractions | Fixed: $M_{i,0} = 1$ | Normalization of tissue states at submersion |
| $w_i$ | Anatomical weights in microbial growth driver $\Phi_M$ | Fixed: $w_H = w_T = w_L = 1/3$ | No region-specific microbial growth data; uniform weighting avoids unidentifiable parameter combinations |
| $\beta_i$ | Anatomical weights in oxygen demand $\Psi_M$ | Fixed: $\beta_H = \beta_T = \beta_L = 1/3$ | Regional oxygen consumption not observable; uniform weighting minimizes assumptions |
| $\lambda_{i,0}$<br>($i = H, T, L$) | Baseline tissue degradation coefficient for anatomical region $i$ | Fixed: $\lambda_{H,0} = \lambda_{T,0} = \lambda_{L,0} = 1$ | No region-specific tissue degradation data are available; equal baseline coefficients provide a neutral reference parameterization and avoid introducing unsupported regional differences |
| $\alpha_{B,i}$<br>($i = H, T, L$) | Regional weighting of microbial contribution to tissue loss | Fixed: $\alpha_{B,H} = \alpha_{B,T} = \alpha_{B,L} = 1$ | No region-specific data are available to distinguish microbial effects among anatomical regions; equal weights avoid introducing unsupported regional heterogeneity |
| $B_{\max}$ | Maximum normalized microbial activity | Fixed: $B_{\max} = 1$ | $B(t)$ is a dimensionless, normalized activity variable; fixing the scale prevents confounding with growth parameters |
| $B_0$ | Initial microbial activity | Fixed: $B_0 = 10^{-3}$ | Small but non-zero seed required for logistic growth; does not affect long-term dynamics |
| $\chi_S$ | Scavenger module switch | Fixed: $\chi_S = 0$ (Palazzo calibration) | Negligible scavenging activity reported for Northern Adriatic Sea cases |
| $\chi_O$ | Oxygen module switch | Fixed: $\chi_O = 1$ | Oxygen effects included through prescribed environmental forcing |
| **(ii) Environmental forcings (prescribed inputs)** |  |  |  |
| $T_{\text{env}}(t)$ | Water temperature forcing | Prescribed time-dependent profile, typically 8–24°C (Northern Adriatic) | Case-specific environmental records as in Palazzo et al. (2020); not estimated from decomposition data |
| $O_{\infty}(t)$ | Bulk dissolved oxygen concentration | Prescribed constant or seasonal average, typically 6–9 mg L<sup>-1</sup> | Independent oceanographic measurements for the Northern Adriatic Sea; treated as external forcing |
| **(iii) Effective parameters estimated from data** |  |  |  |
| $k_{B,0}$ | Baseline microbial attack coefficient | 1.91 | This study |
| $\eta_B$ | Microbial growth efficiency (effective) | 0.05 | This study |
| $\mu_{B,0}$ | Baseline microbial loss rate | 1.21 | This study |
| $c_B$ | Oxygen consumption conversion factor | 38.5 | This study |
| $\phi$ | Near-body oxygen renewal rate | 0.156 | This study |
| $b$ | Tissue-loss-to-score scaling factor | 29.6 | This study |

##### Tissue-related parameters

- *λ*_*i*,0_: baseline tissue degradation rate for anatomical region *i* ∈{*H, T, L*} under reference environmental conditions.
- *Q*_10,*i*_: temperature sensitivity coefficient for tissue degradation in region *i*, representing the multiplicative change in degradation rate for a 10^°^C increase in temperature.
- *M*_*i*,0_: initial normalized tissue mass fraction for region *i* at the time of submersion.

##### Microbial activity parameters

- *B*(*t*): normalized microbial activity associated with the body.
- *k*_*B*,0_: baseline microbial attack coefficient at the reference temperature.
- *Q*_10,*B*_: temperature sensitivity of microbial activity.
- *η*_*B*_: microbial growth efficiency coefficient.
- *B*_max_: effective carrying capacity for microbial activity.
- *µ*_*B*,0_: baseline microbial loss rate at the reference temperature.
- *Q*_10,*µ*_: temperature sensitivity of the microbial loss rate.
- *α*_*B,i*_: weighting coefficient representing the contribution of microbial activity to tissue loss in region *i*.

##### Oxygen-related parameters

- *O*_*s*_(*t*): dissolved oxygen concentration in the near-body water layer.
- *O*_∞_(*t*): dissolved oxygen concentration in the surrounding bulk water.
- *K*_*O*_: half-saturation constant controlling the strength of oxygen limitation on tissue degradation.
- *ϕ*: exchange or renewal rate between near-body water and the bulk water, representing mixing, diffusion and small-scale flow.
- *c*_*B*_: conversion coefficient linking microbial activity and tissue availability to oxygen consumption.
- *β*_*i*_: weighting coefficient for the contribution of tissue region *i* to oxygen demand.

##### Scavenger-related parameters (optional module)

- *σ*_scav,*i*_: effective scavenger pressure acting on tissue region *i*, representing the average impact of fish and other macro-fauna.
- *σ*_*O*_: effective background oxygen demand due to scavenger activity and associated fauna.

##### Environmental and structural parameters

- *T*_env_(*t*): ambient water temperature as a function of time.
- *T*_ref_ : reference temperature used for defining temperature-dependent rates.
- *w*_*i*_: anatomical weighting coefficients satisfying ∑_*i*_ *w*_*i*_ = 1, used to aggregate tissue availability for microbial growth.
- *χ*_*S*_: binary switch activating (*χ*_*S*_ = 1) or deactivating (*χ*_*S*_ = 0) scavenger effects.
- *χ*_*O*_: binary switch activating (*χ*_*O*_ = 1) or deactivating (*χ*_*O*_ = 0) oxygen limitation.

#### 3.1.2 Tissue dynamics

For each anatomical region *i* ∈ *{H, T, L*} (head/neck, trunk, limbs), the temperature- and (optionally) oxygen-modulated tissue degradation rate is defined as

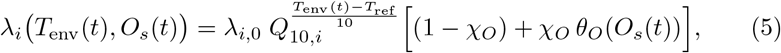

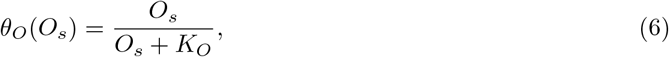

where *λ*_*i*,0_ *>* 0, *Q*_10,*i*_ *>* 0, *T*_ref_ and *K*_*O*_ *>* 0 are fixed parameters. The temperature dependence follows a Q10-type exponential scaling, a standard form in metabolic theory describing how biological rates approximately double for every 10°C rise in temperature. Oxygen limitation is modeled via a Michaelis-Menten-type saturation function, whereby degradation is strongly suppressed at low oxygen and approaches its unconstrained rate as oxygen becomes abundant.

Tissue degradation is assumed to result from the interaction between the remaining degradable tissue and the local microbial activity. Since decomposition requires both remaining degradable tissue and microbial activity, the effective tissue-loss rate is modeled using a mass-action-type interaction proportional to *B*(*t*)*M*_*i*_(*t*). This bilinear formulation represents the simplest reduced macroscopic description of microbial decomposition, capturing the aggregate effect of microbial activity on tissue loss while preserving biological interpretability. Temperature and oxygen availability modulate this baseline interaction through the factors *k*_*B*_(*T*_env_) and *λ*_*i*_(*T*_env_, *O*_*s*_). The objective of the present model is not to describe the underlying biochemical mechanisms in detail, but rather to represent their overall effect on tissue degradation at the macroscopic scale. Microbial activity and (optional) scavenger pressure contribute additively to tissue loss. The tissue dynamics are given by

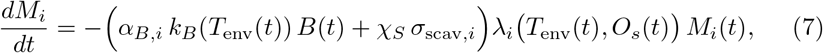

for *i* ∈ *{H, T, L}*, where *α*_*B,i*_ ≥ 0 weights the contribution of microbial activity and *σ*_scav,*i*_ ≥ 0 represents an effective regional scavenger pressure, active only when *χ*_*S*_ = 1.

The temperature dependence of microbial attack is modelled as

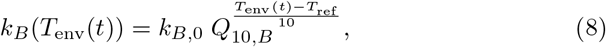

with baseline coefficient *k*_*B*,0_ *>* 0 and microbial *Q*_10,*B*_ *>* 0.

#### 3.1.3 Microbial activity dynamics

Microbial activity *B*(*t*) is described by a logistic-type equation driven by available tissue:

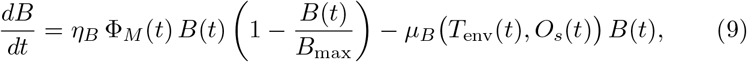

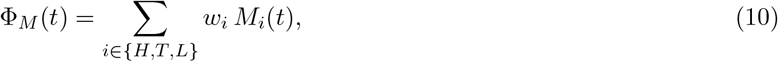

where *w*_*i*_ *>* 0 with ∑_*i*_ *w*_*i*_ = 1, *η*_*B*_ *>* 0 is a growth efficiency and *B*_max_ *>* 0 is an effective carrying capacity.

The microbial loss rate is defined as

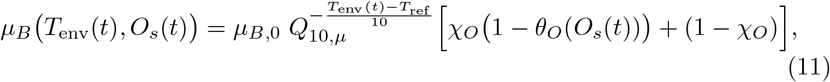

so that oxygen dependence is removed when *χ*_*O*_ = 0.

#### 3.1.4 Near-body oxygen dynamics

When *χ*_*O*_ = 1, the near-body oxygen concentration follows the box model

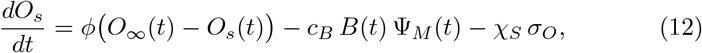

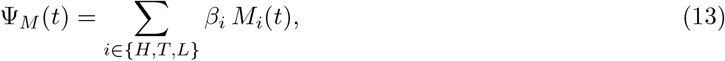

where *ϕ >* 0 is the exchange rate with the bulk water, *c*_*B*_ *>* 0 converts microbial activity into oxygen consumption, *β*_*i*_ ≥ 0 are regional weights, and *σ*_*O*_ ≥ 0

represents an effective background oxygen demand due to scavengers when *χ*_*S*_ = 1.

When *χ*_*O*_ = 0, we assume

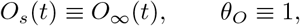

and equation (12) is not integrated.

#### 3.1.5 Initial conditions

At the time of submersion (*t* = 0) we set

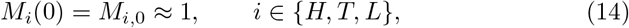

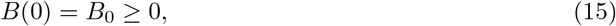

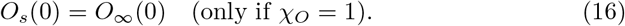

## 4 Calibration using the Palazzo dataset

Given the absence of publicly available case-level datasets, the published TADS–PMSI relationship provides the most comprehensive quantitative reference currently available for calibration in the Northern Adriatic setting.It is important to clarify the nature of the calibration performed in this study. Rather than fitting the mechanistic model directly to individual case-level time series, we calibrate the model to reproduce the published empirical relationship between total aquatic decomposition score (TADS) and postmortem submersion interval (PMSI) reported by Palazzo et al. for Northern Adriatic Sea cases [4]. This regression provides an aggregated description of score progression in time, which we treat as a macroscopic target for the underlying biological dynamics.

Accordingly, the mechanistic ODE system is calibrated so that its predicted score trajectory TADS(*t*) follows the published regression over a specified PMSI range used for fitting. In this sense, the empirical regression is not replaced but embedded as a calibration target within a mechanistic framework, allowing inference of effective (environment- and dataset-specific) parameters governing tissue degradation, microbial activity, and oxygen limitation. Once calibrated over the fitted range, the resulting monotone score–time relationship provides a well-defined inverse mapping from observed decomposition scores to PMSI estimates within that same range and under the assumed environmental forcings.

For the Northern Adriatic Sea cases reported by Palazzo et al. [6], negligible scavenging activity was described. Accordingly, in the calibration presented here, we fix:S

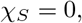

and exclude all scavenger-related parameters from estimation.

Although case-specific dissolved oxygen measurements are not available in the Palazzo dataset, bulk dissolved oxygen levels for the Northern Adriatic Sea are documented in independent oceanographic studies (e.g. [14]). We therefore activate the oxygen module by fixing

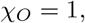

and prescribe the bulk oxygen concentration *O*_∞_(*t*) as an external environmental forcing (constant or seasonal-average profile as specified below), while estimating effective oxygen-related exchange/consumption parameters from decomposition score data.

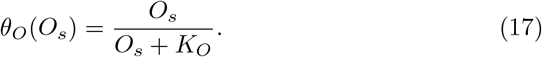

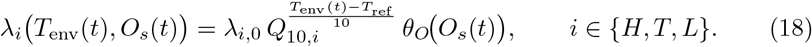

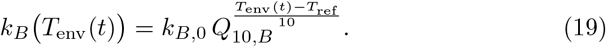

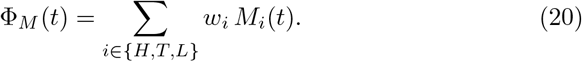

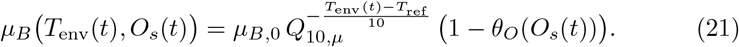

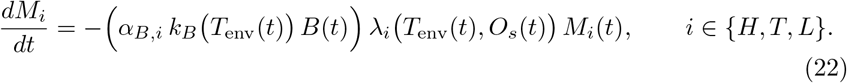

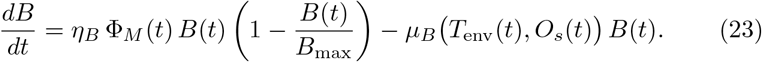

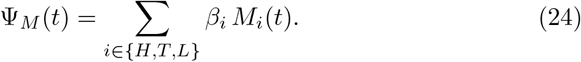

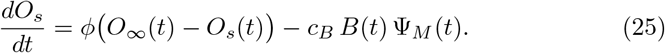

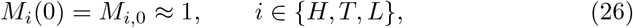

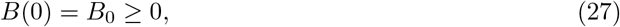

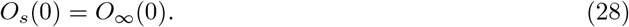

The reduced parameter vector to be estimated is

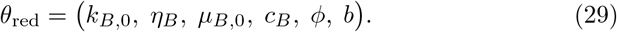

### 4.1 Observation model: mapping tissue loss to decomposition scores

Regional decomposition scores were computed using the observation model defined in the Methods section (Eqs. (1)–(3)). The intercept of each regional score was fixed at 1, while a common slope parameter was estimated across all anatomical regions, yielding the model-predicted FAD, BAD, LAD, and TADS values.

### 4.2 Calibration target and objective function (aggregated regression)

Let TADS^reg^(*t*) denote the published regression function from [4]. Since the Total Aquatic Decomposition Score (TADS) is defined as the sum of the three regional decomposition scores (FAD, BAD, and LAD), each having a minimum possible score of 1, the observation-model intercept was fixed at *a* = 3. Consequently, we calibrate the reduced parameter vector

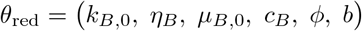

by minimizing the squared deviation between the mechanistic score trajectory and the regression target over a grid of times 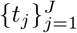 spanning the fitted PMSI range:

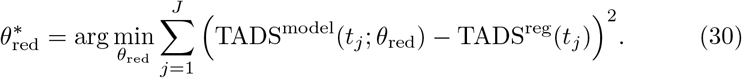

Details of the time grid, numerical solver, and optimization routine are provided in the Methods section.

### 4.3 Inverse use and scope of applicability

Within the fitted PMSI range and under the assumed environmental forcings, the monotonicity of TADS^model^(*t*) allows inversion to obtain PMSI estimates from observed TADS values. The present proof-of-concept calibration is specific to the Northern Adriatic context represented by the published regression and the prescribed environmental forcings; application to other aquatic settings would require comparable environmental characterization and re-calibration using appropriate reference data.

### 4.4 Case-level calibration as future work

When case series with medico-legally established PMSI values and corresponding decomposition scores are available, the same model structure can be calibrated directly at the case level by integrating the ODE system to each case’s recovery time and fitting to observed scores. This extension is not pursued here due to limitations in publicly available case-level PMSI reporting.

The proposed calibration framework jointly estimates the effective parameters of the mechanistic ODE model and the observation-model slope parameter that maps latent tissue loss to observed decomposition scores. In this way, the decomposition scores reported by Palazzo et al. are used both to constrain the underlying biological dynamics and to estimate the observation-model scaling parameter relating latent tissue loss to decomposition scores, yielding a monotone and invertible relationship between decomposition scores and postmortem interval.

Although the estimated parameters should not be interpreted as uniquely identifiable biological rate constants, their relative magnitudes remain biologically plausible within the context of the reduced model. In particular, the relatively small microbial growth efficiency (*η*_*B*_), together with the larger microbial loss coefficient (*µ*_*B*,0_), produces bounded microbial activity throughout the calibrated interval, while the estimated oxygen-consumption coefficient (*c*_*B*_) is consistent with substantial coupling between microbial activity and local oxygen depletion. These observations provide qualitative support for the biological consistency of the calibrated effective parameter set.

Before presenting the calibration results, we briefly summarize the role of the empirical regression reported by Palazzo et al. in our modeling framework. That regression provides a widely used relationship between postmortem interval (PMSI) and the total body decomposition score (TADS), but does not explicitly encode the underlying biological processes driving tissue degradation. In our approach, this empirical relationship serves as a calibration target for the mechanistic model, allowing us to anchor the latent tissue degradation dynamics to observed forensic scoring data. Figure 2 illustrates the resulting fit of the mechanistic score trajectory to the published regression, which forms the basis for the inverse PMSI estimation discussed subsequently.

**Figure 2.**
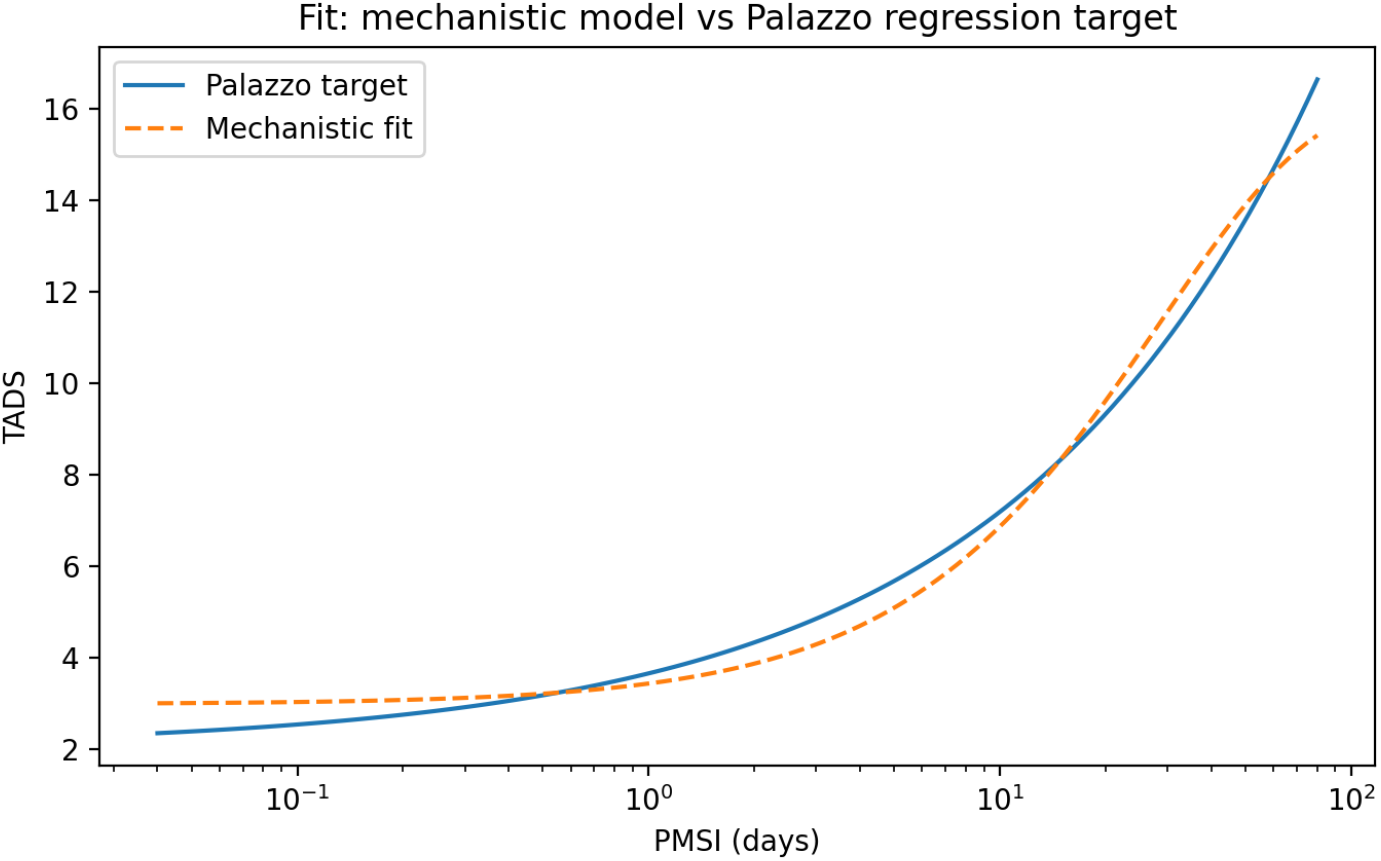
Calibration of the mechanistic model to the Palazzo TADS– PMSI regression. The solid curve shows the published regression target TADS = 2.01908 + 1.63397 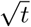 with *t* in days. The dashed curve is the mechanistic model prediction TADS(*t*) obtained by least-squares fitting over *t*∈ [0.04, 80] days (log-spaced grid).

To complement the visual comparison shown in Figure 2, the quality of the fit was further quantified numerically. Over the fitted log-spaced time grid (*n* = 120 points, *t* ∈ [0.04, 80] days), the calibrated mechanistic trajectory reproduces the Palazzo regression target with a coefficient of determination of *R*^2^ = 0.986 (adjusted *R*^2^ = 0.986, accounting for the six fitted parameters), a root-mean-square error of RMSE = 0.44 TADS units, and a mean absolute error of MAE = 0.39 TADS units. The Pearson correlation coefficient between the model trajectory and the regression target was *r* = 0.993. The maximum absolute deviation (1.22 TADS units) occurred at the upper boundary of the fitted interval (*t* = 80 days), indicating a modest systematic divergence at the longest postmortem submersion intervals considered, consistent with the slight departure between the fitted and target curves visible at the right-hand end of Figure 2. Table 2 summarizes these goodness-of-fit metrics.

**Table 2:** Goodness-of-fit metrics for the calibrated mechanistic model relative to the Palazzo regression target (Figure 2).

| **Metric** | **Value** |
| --- | --- |
| Number of grid points ($n$) | 120 |
| Number of fitted parameters ($p$) | 6 |
| RMSE | 0.44 |
| MAE | 0.39 |
| MAPE | 9.14% |
| $R^2$ | 0.986 |
| Adjusted $R^2$ | 0.986 |
| Pearson correlation ($r$) | 0.993 |
| Maximum absolute error (at $t$) | 1.22 (at $t = 80$ days) |

Having established a calibrated mechanistic relationship between decomposition score and postmortem interval, we next consider its inverse use. In forensic practice, the observed quantity is typically a decomposition score, whereas the postmortem interval is unknown. The monotonicity of the fitted mechanistic score trajectory, therefore, enables inversion, allowing the postmortem interval to be estimated directly from an observed TADS value. In Figure. 3, we illustrate this inverse mapping by computing PMSI estimates for the example cases reported by Palazzo, based on their published decomposition scores.

**Figure 3.**
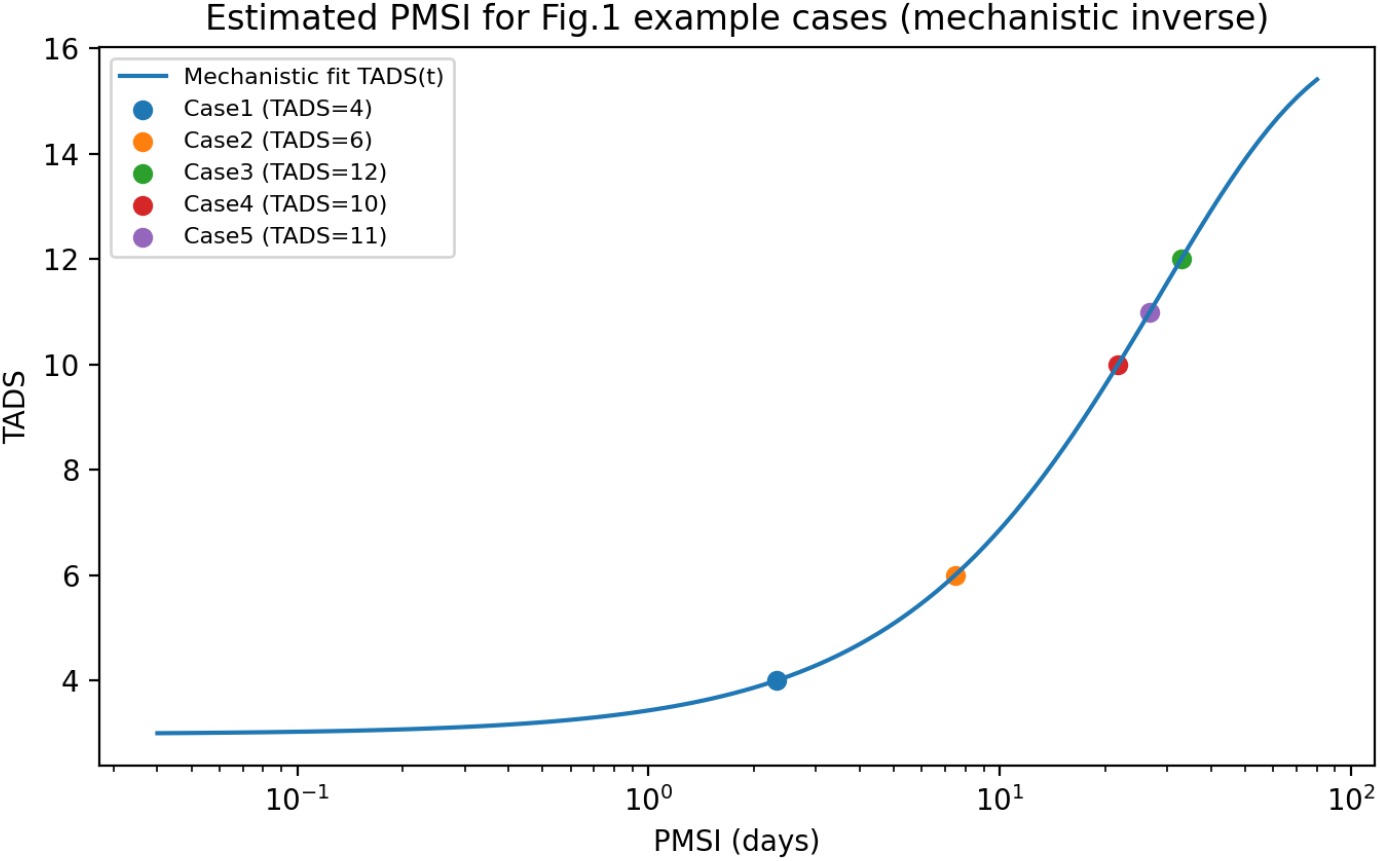
Mechanistic inverse mapping from TADS to PMSI for the Fig. 1 example cases. Using the fitted mechanistic curve TADS(*t*) (blue), we compute PMSI estimates by inversion, i.e., finding *t* such that TADS(*t*) = TADS_obs_. Colored markers indicate the estimated PMSI values corresponding to the example cases reported by Palazzo et al. (Case1–Case5), based on their published TADS values. The monotonicity of TADS(*t*) over the fitted range ensures a unique inverse.

Figures 2 and 3 together summarize the practical outcome of the proposed calibration approach. The fitted mechanistic trajectory reproduces the overall trend of the published TADS–PMSI regression while introducing a biologically motivated, smooth score evolution.Importantly, the resulting score–time relationship remains monotone over the fitted range, enabling a consistent inverse mapping from observed decomposition scores to postmortem interval estimates. The example cases illustrate that the mechanistic inverse yields PMSI values comparable to those obtained from the empirical regression, while providing a process-based interpretation of score progression.

## 5 Observation Strategies for Mechanistic Model Calibration

### 5.1 Motivation and Observation Design

In the previous section, the proposed mechanistic model was successfully calibrated against the Total Aquatic Decomposition Score (TADS) data reported by Palazzo et al. [4]. The resulting agreement demonstrated that the model is capable of reproducing the overall decomposition trajectory observed in submerged remains. However, obtaining an accurate fit using TADS observations alone does not necessarily imply that the underlying biological parameters are uniquely or reliably determined. Since TADS provides only an aggregate description of decomposition progression, different parameter combinations may produce nearly indistinguishable decomposition trajectories while corresponding to substantially different underlying biological processes. This naturally raises the question of whether incorporating additional experimentally accessible observations can improve parameter estimation in mechanistic decomposition models.

Recent developments in aquatic forensic science provide a strong motivation for addressing this question. While decomposition scoring remains the primary source of information for PMSI estimation, several studies have explored complementary biological and environmental observations. Wallace et al. [18] demonstrated that microbial communities on submerged carcasses undergo reproducible successional changes during decomposition, suggesting that microbial observations contain valuable temporal information. Building upon this idea, Jangid et al. [19] showed that microbial community composition can be exploited for PMSI estimation in human remains using machine-learning techniques. In addition to microbial observations, Bone et al. [20] investigated the relationships between microbial communities and environmental variables, including dissolved oxygen, during aquatic decomposition experiments. More recently, Bray et al. [21] examined several water-quality measurements as potential indicators of PMSI. Collectively, these studies illustrate the growing interest in observations that capture biological and physicochemical processes beyond external decomposition scores.

The mechanistic model proposed in this study naturally incorporates two state variables closely related to these emerging observation types: microbial activity (*B*) and dissolved oxygen concentration (*O*). Although microbial activity is represented as an aggregate biological process rather than a directly measurable quantity, experimentally accessible observations such as microbial community composition or bacterial abundance obtained through molecular techniques provide biologically related information. Likewise, dissolved oxygen is an experimentally measurable environmental variable that has already been incorporated into recent aquatic decomposition studies. Consequently, the proposed model provides a natural framework for investigating how different combinations of experimentally accessible observations influence parameter calibration.

To address this question, we performed a series of synthetic observation experiments using the calibrated parameter set obtained in the previous section as the ground-truth parameter vector. The mechanistic model was simulated to generate reference trajectories for decomposition score, microbial activity, and dissolved oxygen concentration, after which measurement noise was added to produce synthetic observations. Four observation scenarios were subsequently considered: (i) decomposition score only (TADS), (ii) TADS together with microbial observations, (iii) TADS together with dissolved oxygen observations, and (iv) the combination of TADS, microbial observations, and dissolved oxygen observations. All parameter estimation procedures were performed using identical optimization algorithms, initial parameter guesses, and parameter bounds so that differences in estimation performance could be attributed solely to the available observation strategy.

### 5.2 Synthetic Observation Generation

To evaluate the contribution of different observation strategies under controlled conditions, synthetic observations were generated from the calibrated mechanistic model rather than using experimental datasets. Although decomposition score, microbial measurements, and water-quality variables have all been reported in the literature, no single experimental dataset provides the combination of observations required for the comparative calibration framework considered in this study. Generating synthetic observations from the calibrated model therefore enables each observation strategy to be evaluated under identical conditions, ensuring that differences in parameter estimation arise solely from the available observations rather than from variations in experimental settings or measurement protocols.

The calibrated parameter set obtained in Section 4 was adopted as the ground-truth parameter vector throughout this study. Using these parameter values, the mechanistic model was solved numerically to generate reference trajectories for all model state variables over the observation period. In particular, the decomposition score, microbial activity, and dissolved oxygen concentration were extracted as the observable quantities considered in the subsequent calibration experiments.

To emulate realistic experimental measurements, synthetic observations were generated by sampling the reference trajectories at the prescribed observation times and adding random measurement noise. The observation times were chosen to be consistent with the temporal sampling adopted during model calibration, while the added noise represents unavoidable experimental uncertainty associated with biological and environmental measurements. Consequently, each synthetic dataset consists of noisy observations of the underlying model trajectories rather than exact model solutions. An example of the resulting synthetic microbial observations is shown in Figure 4.

**Figure 4.**
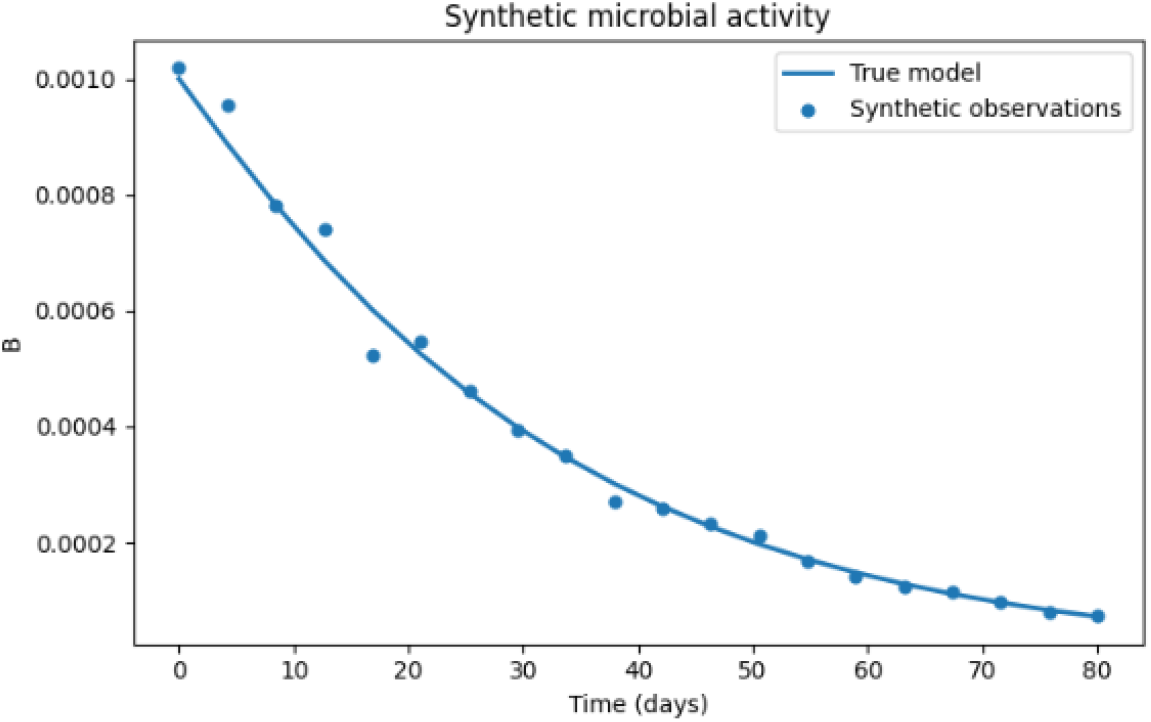
Generation of synthetic microbial observations from the calibrated mechanistic model. The solid curve represents the ground-truth microbial activity trajectory obtained using the calibrated parameter set, while the circular markers denote synthetic observations generated by sampling the reference trajectory at the prescribed observation times and adding random measurement noise. Although only microbial activity is illustrated, the same synthetic observation generation procedure was applied to all observable quantities considered in this study.

Based on these synthetic observations, four different observation scenarios were constructed to investigate how the availability of additional measurements influences parameter estimation. The first scenario used decomposition score (TADS) observations alone, representing the conventional calibration strategy based solely on external decomposition scoring. The second scenario combined TADS observations with microbial observations, whereas the third combined TADS observations with dissolved oxygen measurements. Finally, the fourth scenario incorporated all three observation types simultaneously, representing the most informative observation strategy considered in this study.

The resulting synthetic datasets formed the basis for all subsequent calibration analyses. In the following sections, parameter estimation was performed independently for each observation scenario, enabling a systematic comparison of how different observation strategies influence parameter recovery and the reliability of the estimated model parameters.

### 5.3 Influence of Observation Strategies on Parameter Recovery and PracticalIdentifiability

The four observation scenarios were compared to investigate how different biological measurements influence both the predictive performance and the practical identifiability of the proposed mechanistic decomposition model. Model performance was first evaluated by comparing the reconstructed trajectories with the synthetic observations, whereas practical identifiability was subsequently assessed using profile likelihood analysis.

Although all four observation strategies reproduced the decomposition score with nearly identical accuracy, their ability to reconstruct the underlying biological variables differed substantially. The TADS root mean squared error (RMSE) remained nearly unchanged across all observation scenarios, ranging from 0.266 to 0.269. The coefficient of determination was also consistently high, with *R*^2^ = 0.9952 for the TADS-only calibration and *R*^2^ = 0.9951 for the TADS+*B* calibration. These results indicate that decomposition score observations can be fitted extremely well under multiple parameter combinations and therefore do not, by themselves, guarantee accurate reconstruction of the underlying biological processes.

The differences between the observation strategies became more evident when microbial activity was examined. As shown in Figure 5(a), calibration using only TADS observations reconstructed microbial activity with a relative RMSE of 10.39%. Incorporating direct microbial observations reduced this error to 5.29%, corresponding to an approximately two-fold improvement in microbial trajectory reconstruction. Interestingly, dissolved oxygen observations also improved the microbial activity prediction even though microbial observations were not included in the calibration. Under the TADS+*O* strategy, the microbial relative RMSE decreased to 8.75%, as illustrated in Figure 5(b). When microbial and oxygen observations were used simultaneously, the microbial relative RMSE was 5.35%, which was nearly identical to that obtained using microbial observations alone. This suggests that direct microbial measurements contain most of the information required to reconstruct the microbial trajectory, while oxygen observations provide a smaller but measurable indirect contribution.

**Figure 5.**
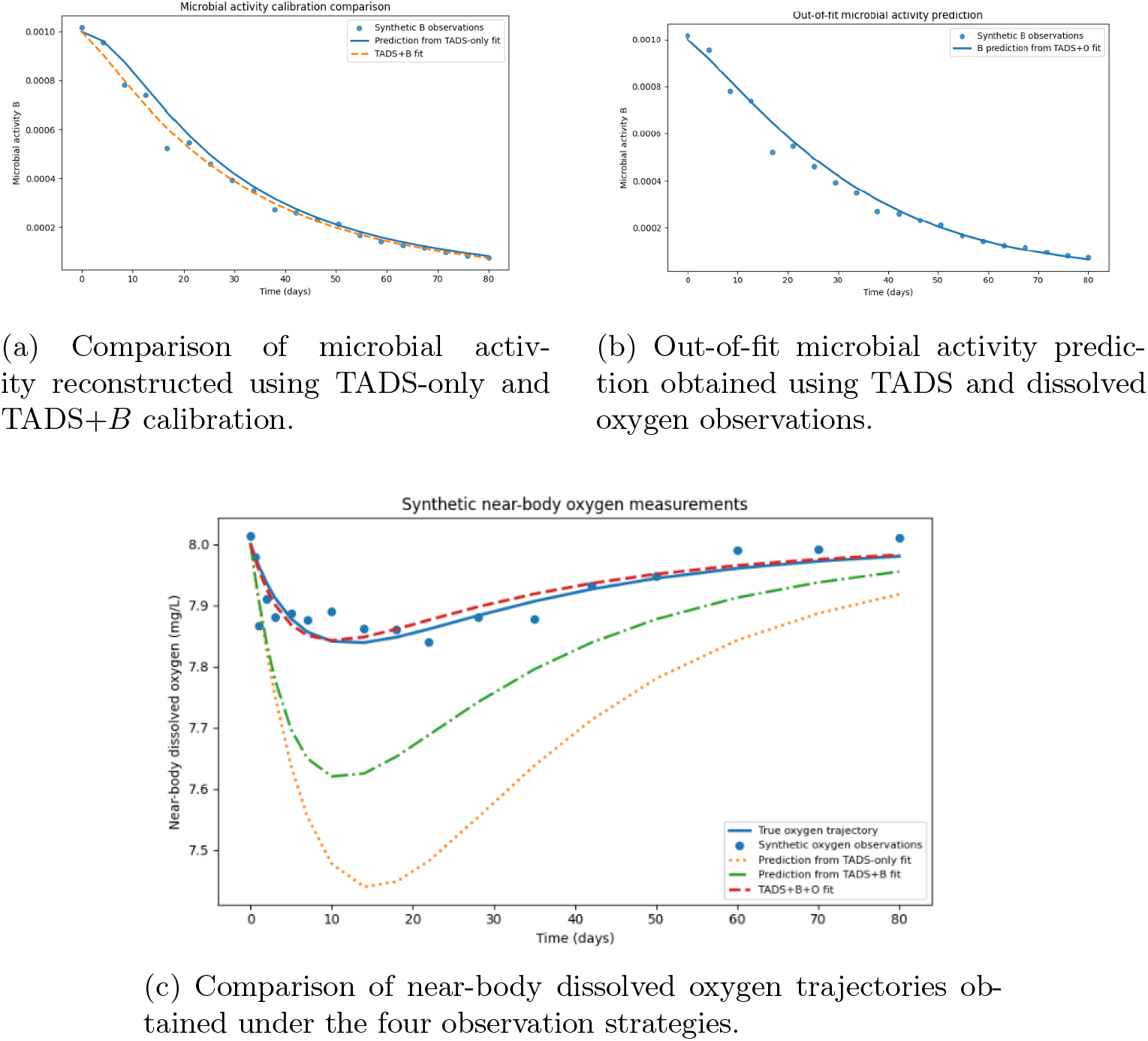
Influence of the observation strategy on the reconstruction of microbial activity and dissolved oxygen. Panel (a) compares microbial activity predictions obtained from TADS-only and TADS+*B* calibration. Panel (b) shows the microbial activity predicted using TADS and dissolved oxygen observations, without directly including microbial measurements in the calibration. Panel (c) compares dissolved oxygen predictions obtained using TADS-only, TADS+*B*, TADS+*O*, and TADS+*B* + *O* observation strategies. Symbols represent synthetic observations, while curves represent model predictions obtained using the estimated parameter sets.

A complementary pattern was observed for dissolved oxygen. Figure 5(c) compares oxygen predictions obtained under the four observation strategies. When only TADS observations were used, the oxygen trajectory was reconstructed with an RMSE of 0.249 mg*/*L. The inclusion of microbial observations reduced the oxygen RMSE to 0.140 mg*/*L, reflecting the mechanistic coupling between microbial activity and oxygen consumption. The greatest improvement occurred when dissolved oxygen observations were included directly in the calibration. The oxygen RMSE decreased to 0.0311 mg*/*L under the TADS+*O* strategy and to 0.0314 mg*/*L under the TADS+*B* + *O* strategy. Thus, microbial measurements improve oxygen prediction indirectly, whereas direct oxygen observations are required for accurate reconstruction of the oxygen dynamics.

Improved trajectory reconstruction does not necessarily imply that the underlying parameters are uniquely constrained. This distinction is revealed by the profile likelihood analysis shown in Figure 6. For the microbial growth coefficient *k*_*B*,0_, the likelihood profile remained open on the upper side when only TADS observations were used. After microbial observations were incorporated, the profile became fully closed, with an approximate 95% confidence interval of 0.897–2.437 under the TADS+*B* strategy. The corresponding interval under the TADS+*B* + *O* strategy was 0.909–2.636. The limited additional narrowing produced by oxygen observations indicates that microbial measurements constitute the principal source of information for constraining *k*_*B*,0_.

**Figure 6.**
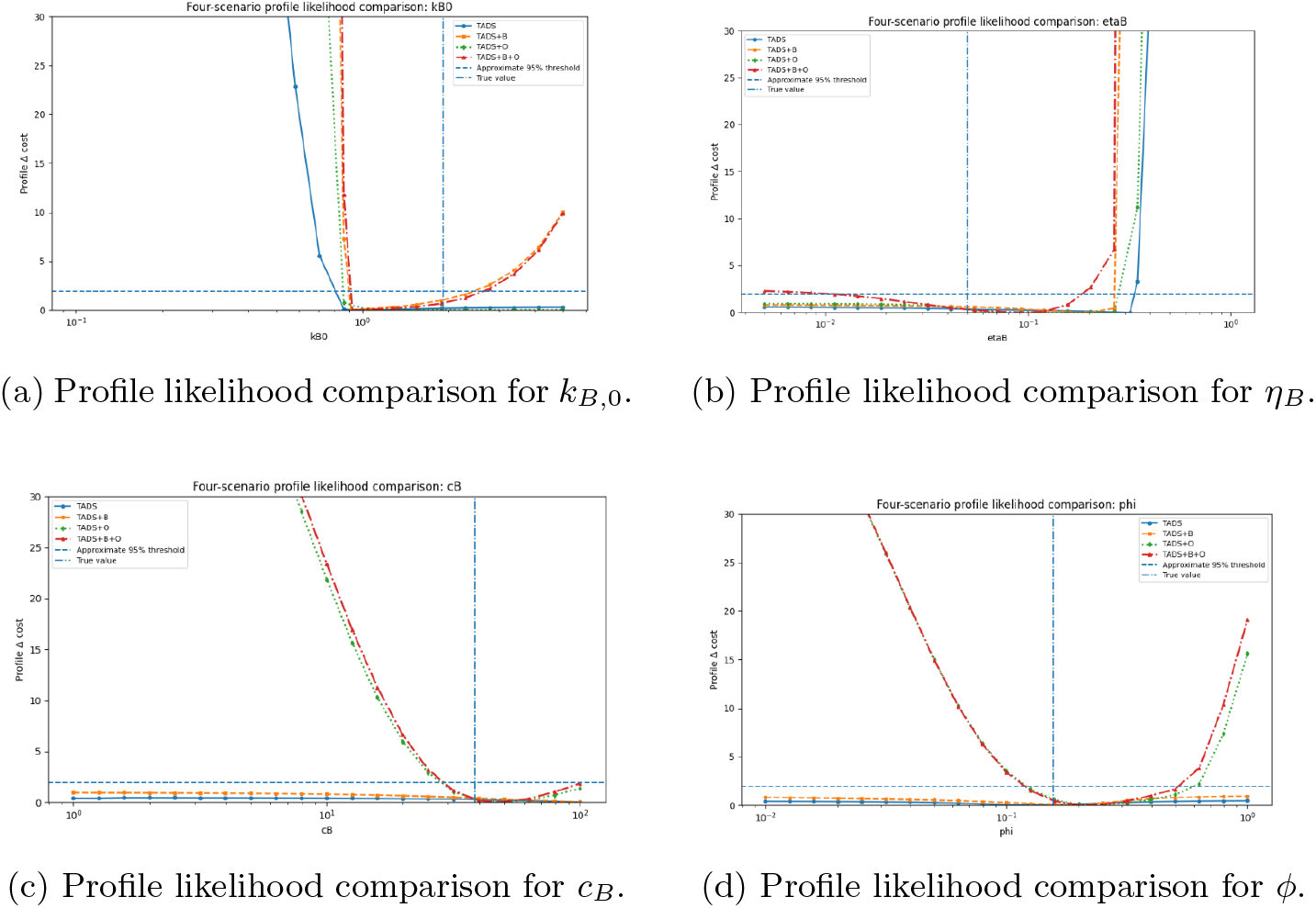
Profile likelihood comparison across the four observation strategies for selected model parameters. The horizontal dashed line denotes the approximate 95% likelihood threshold, while the vertical dash-dotted line indicates the ground-truth parameter value used to generate the synthetic observations. Closed profiles indicate that both lower and upper confidence limits can be determined, whereas open profiles indicate that the corresponding parameter remains only partially constrained by the available observations.

A different pattern was observed for *η*_*B*_.Its profile remained open on the lower side under the TADS-only, TADS+*B*, and TADS+*O* strategies. A fully closed profile was obtained only when microbial and dissolved oxygen observations were combined, yielding an approximate 95% confidence interval of 0.0113–0.1834. This result indicates that neither observation type alone contains sufficient information to constrain *η*_*B*_, whereas the combined data provide complementary information about the interaction between microbial growth and oxygen-dependent decomposition.

A similar behavior was observed for the microbial loss parameter *µ*_*B*,0_. As with *η*_*B*_, the likelihood profile remained open when either microbial activity or dissolved oxygen observations were considered separately and became closed only under the combined TADS+*B* + *O* observation strategy. This finding indicates that reliable estimation of the microbial loss rate also requires complementary information from both microbial and oxygen dynamics, reflecting the intrinsic coupling between microbial growth and decay processes within the model.

The oxygen-related parameters exhibited similarly distinct responses to the observation design. For *c*_*B*_, both the lower and upper sides of the likelihood profile remained open when only TADS or TADS+*B* observations were used. The inclusion of dissolved oxygen observations strongly constrained the lower side of the profile, producing lower confidence limits of approximately 28.37 and 29.02 under the TADS+*O* and TADS+*B* + *O* strategies, respectively. However, the upper confidence bound remained open in both cases, indicating that *c*_*B*_ became partially, but not fully, identifiable.

In contrast, the oxygen replenishment parameter *ϕ* became fully identifiable once dissolved oxygen observations were included.Its profile was open on both sides under the TADS-only and TADS+*B* strategies but closed under both oxygen-containing observation designs. The approximate 95% confidence interval was 0.121–0.597 for TADS+*O* and 0.119–0.515 for TADS+*B* + *O*. These results demonstrate that dissolved oxygen observations provide the primary information required to constrain the oxygen replenishment process represented by *ϕ*.

Overall, the results demonstrate that different observation types contribute complementary information to the calibration process. Direct microbial observations primarily improve the reconstruction of microbial activity and constrain the microbial growth coefficient *k*_*B*,0_. Dissolved oxygen observations provide the strongest information for the oxygen-consumption parameter *c*_*B*_ and the replenishment parameter *ϕ*, while the combined use of microbial and oxygen measurements is required to close the likelihood profile of *η*_*B*_. Although TADS observations alone are sufficient to reproduce the decomposition score with high accuracy, they are insufficient to uniquely determine several mechanistic parameters. The incorporation of biologically informative observations therefore improves both the predictive interpretation and the practical identifiability of the proposed decomposition model.

Finally, the mapping parameter *b* remained practically non-identifiable under all four observation strategies. Unlike the mechanistic parameters governing the decomposition dynamics, *b* belongs to the observation model that maps the underlying tissue-state variables to TADS. Because the observation model parameters *a* and *b* exhibit strong compensation, simultaneous estimation resulted in an ill-conditioned calibration problem, and *a* was therefore fixed at 3. Nevertheless, the profile likelihood of *b* remained open in every observation scenario, indicating that the available observations do not provide sufficient independent information to uniquely constrain the TADS mapping. This result suggests that the remaining uncertainty arises primarily from the observation model rather than from the underlying decomposition dynamics.

### 5.4 Monte Carlo robustness analysis

To determine whether the conclusions obtained from a single synthetic dataset remained valid under different realizations of measurement noise, a Monte Carlo robustness analysis was performed. Twenty independent noisy datasets were generated by perturbing the reference trajectories with newly sampled Gaussian measurement noise. Each dataset was calibrated independently under the four observation strategies, namely TADS, TADS+*B*, TADS+*O*, and TADS+*B* +*O*. To reduce the influence of local minima, every calibration was repeated from four different initial parameter guesses. Thus, each observation strategy involved 20 noisy datasets and four optimization starts per dataset, corresponding to 80 optimization runs per strategy and 320 runs across all four strategies. All reported calibrations converged successfully. Robustness was subsequently evaluated from three complementary perspectives: parameter recovery accuracy, boundary-hit frequency, and predictive performance.

The distributions of the absolute percentage recovery errors for four representative parameters are presented in Figure 7. These parameters were selected to illustrate the distinct information supplied by microbial and dissolved oxygen measurements.

**Figure 7.**
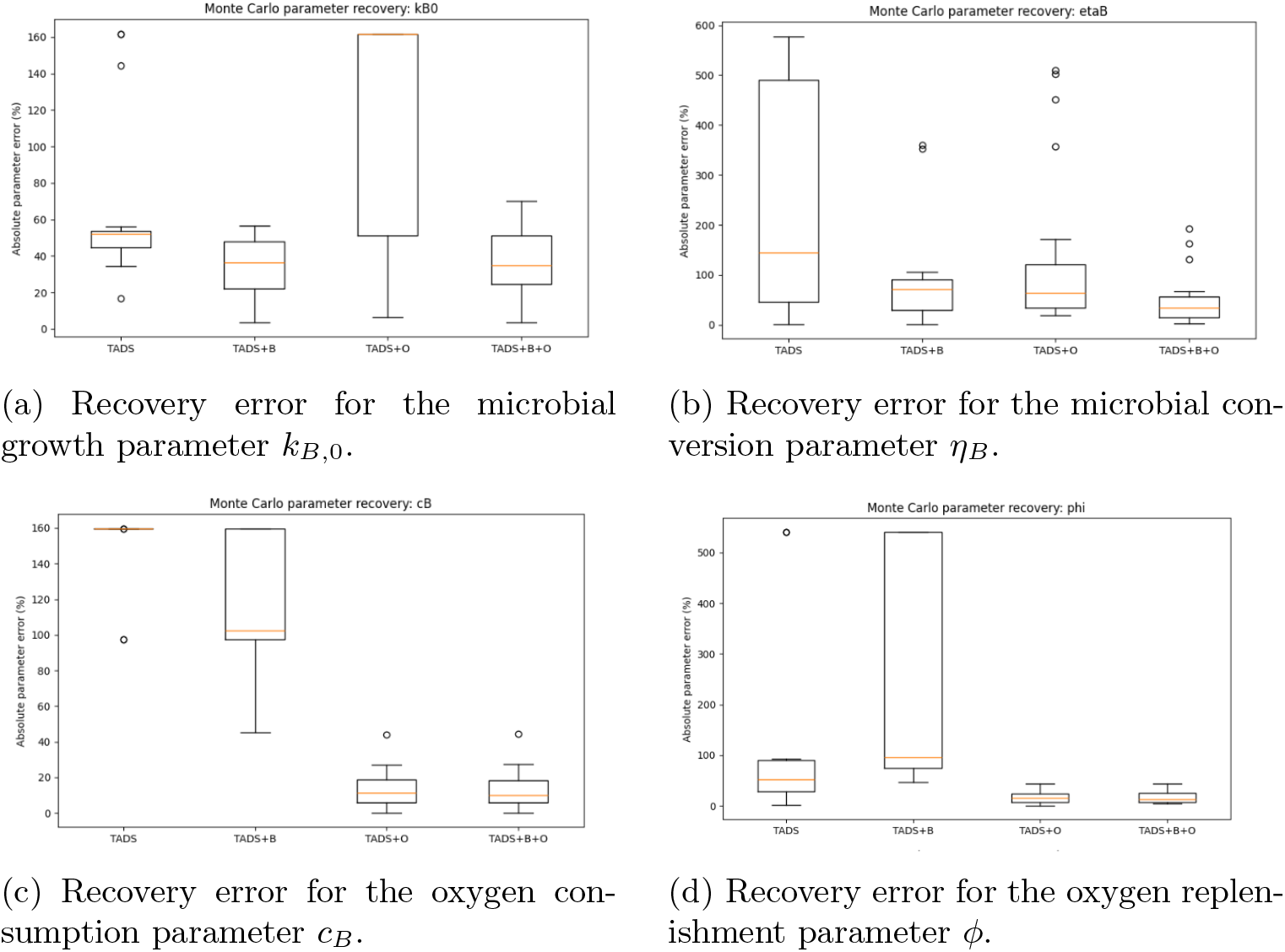
Monte Carlo robustness of parameter recovery across observation strategies. Boxplots show the distributions of absolute percentage recovery errors obtained from 20 independent noisy datasets under the four observation strategies. The panels correspond to (a) the microbial growth parameter *k*_*B*,0_, (b) the microbial conversion parameter *η*_*B*_, (c) the oxygen consumption parameter *c*_*B*_, and (d) the oxygen replenishment parameter *ϕ*. Lower median errors and narrower interquartile ranges indicate more accurate and robust parameter recovery across repeated realizations of measurement noise.

As shown in Figure 7, the Monte Carlo experiments reproduced the same parameter-specific observation hierarchy identified in Section 5.3. Parameters associated primarily with microbial dynamics were generally recovered more accurately when microbial measurements were available, whereas parameters governing the oxygen dynamics exhibited their largest improvements when dissolved oxygen observations were included.

For the microbial growth parameter *k*_*B*,0_, shown in Figure 7a, the median absolute recovery error decreased from 52.1% under the TADS-only strategy to 36.4% under TADS+*B*. The combined TADS+*B* + *O* strategy produced a comparable median error of 35.0%. By contrast, TADS+*O* produced a median error of 161.8%, together with a broad error distribution. Dissolved oxygen measurements alone therefore did not provide sufficient information to recover the microbial growth rate reliably. Interestingly, the TADS+*O* strategy produced poorer recovery of *k*_*B*,0_ than the TADS-only strategy. This likely reflects the indirect relationship between dissolved oxygen observations and the microbial growth coefficient, whereby oxygen measurements constrain the oxygen subsystem without directly informing *k*_*B*,0_, allowing alternative parameter combinations to provide similarly accurate trajectory fits. This effect was largely mitigated when complementary microbial observations were incorporated.

The recovery of *η*_*B*_ displayed a stronger dependence on complementary observations. As illustrated in Figure 7b, the TADS-only strategy produced a median error of 144.7%, with a particularly wide interquartile range of 46.2– 490.5%. The median error decreased to 72.2% under TADS+*B* and 64.5% under TADS+*O*, but the lowest error, 34.0%, was obtained when microbial and oxygen measurements were incorporated together. A similar result was obtained for the microbial loss parameter *µ*_*B*,0_, whose median recovery error decreased from 91.6% under TADS alone to 21.5% under TADS+*B* + *O*. These results suggest that parameters linking microbial activity to the surrounding aquatic environment are most reliably recovered when both components of the model are observed.

The complementary role of oxygen measurements is particularly evident for the oxygen-related parameters. For *c*_*B*_, shown in Figure 7c, the median recovery error was 159.7% under TADS alone and remained high at 102.6% under TADS+*B*. Once dissolved oxygen measurements were incorporated, however, the median error decreased sharply to 11.4% under TADS+*O* and 9.8% under TADS+*B* + *O*. Similarly, for *ϕ* in Figure 7d, the median error decreased from 51.6% under TADS alone to 15.6% under TADS+*O* and 12.6% under the combined strategy. The TADS+*B* strategy performed particularly poorly for *ϕ*, producing a median error of 95.2% and a wide interquartile range of 75.2– 541.0%. Thus, microbial observations without oxygen measurements did not reliably constrain the processes governing oxygen replenishment. Overall, these results show that direct dissolved oxygen measurements are essential for recovering parameters associated with oxygen consumption and replenishment.

The second robustness measure was the boundary-hit frequency, defined as the proportion of Monte Carlo calibrations in which an estimated parameter reached either the lower or upper bound of its prescribed admissible interval. Frequent boundary hits indicate that the available observations do not sufficiently constrain the parameter within the interior of the feasible parameter space and therefore provide an additional indication of limited practical identifiability. For *c*_*B*_, the boundary-hit frequency was 100% under the TADS-only strategy and remained high at 80% under TADS+*B*.It decreased to 0% under both TADS+*O* and TADS+*B*+*O*. For *ϕ*, the corresponding rate was 30% under TADS alone and increased to 60% under TADS+*B*, whereas no boundary hits were observed under either strategy containing dissolved oxygen measurements. These results are consistent with the recovery-error distributions in Figures 7c and 7d and confirm that oxygen measurements stabilize the estimation of both oxygen-related parameters.

The combined strategy also eliminated boundary hits for several parameters involved in microbial dynamics. The boundary-hit frequency for *η*_*B*_ was 15% under TADS, 25% under TADS+*B*, and 20% under TADS+*O*, but decreased to 0% under TADS+*B* + *O*. Similarly, the boundary-hit frequency for *µ*_*B*,0_ decreased from 25% under TADS and 10% under TADS+*O* to 0% under the combined strategy. These findings closely mirror the profile likelihood results reported in Section 5.3 and provide independent evidence that the improvements in practical identifiability persist across repeated noisy datasets.

The third robustness measure was predictive performance. Median TADS RMSE values remained between 0.296 and 0.299 across all four observation strategies, demonstrating that the observed decomposition-score trajectory could be reproduced with comparable accuracy in every case. However, similar TADS fits did not imply equally accurate recovery of the underlying biological dynamics.

The median relative microbial prediction error was 11.19% under TADS alone and decreased to 4.32% under TADS+*B* and 4.35% under TADS+*B* + *O*. In contrast, the TADS+*O* strategy produced a substantially larger microbial error of 21.16%, despite providing an accurate reconstruction of the dissolved oxygen trajectory. Thus, oxygen measurements alone may constrain the oxygen subsystem while still permitting considerable uncertainty in the unobserved microbial dynamics. Conversely, the median oxygen RMSE was 0.2681 mg/L under TADS alone and 0.1021 mg/L under TADS+*B*, but decreased to 0.0260 mg/L under TADS+*O* and 0.0259 mg/L under TADS+*B* + *O*. These results confirm that accurate prediction of each latent biological variable depends primarily on the availability of observations directly related to that variable.

Overall, the Monte Carlo experiments independently reproduced the principal conclusions of the single-dataset and profile likelihood analyses. Microbial observations improved the recovery and prediction of microbial dynamics, dissolved oxygen measurements were essential for constraining the oxygen subsystem, and the combined strategy generally produced the most balanced parameter recovery. The proposed observation framework therefore yields robust and reproducible parameter estimates under repeated measurement uncertainty, increasing confidence in its future application to experimental and forensic datasets.

## 6 Discussion

Estimating the postmortem submersion interval (PMSI) remains one of the most challenging problems in aquatic forensic science because decomposition results from multiple interacting biological, chemical, and environmental processes. Existing approaches based on accumulated degree-days (ADD) or empirical decomposition score regressions have demonstrated considerable practical value; however, they primarily describe statistical associations rather than the biological mechanisms responsible for decomposition. Consequently, their predictive performance may deteriorate when environmental conditions differ from those under which the original models were established. In contrast, the present study introduces a reduced mechanistic ordinary differential equation (ODE) model that explicitly links aquatic decomposition dynamics to microbial activity and dissolved oxygen processes while preserving direct forensic interpretability.

A principal contribution of this work is that the proposed framework combines biological realism with mathematical tractability. Despite its relatively small number of state variables and parameters, the model successfully reproduces the published TADS–PMSI relationship for Northern Adriatic Sea cases while maintaining a strictly monotone score–time relationship. From a forensic perspective, this property is particularly desirable because increasing decomposition scores always correspond to increasing postmortem submersion intervals, thereby avoiding ambiguous inverse PMSI estimates. More importantly, the model provides biologically interpretable parameters that describe microbial growth, oxygen consumption, and environmental renewal processes rather than merely fitting an empirical regression curve.

Beyond reproducing observed decomposition trajectories, the present study also investigated how different observation strategies influence parameter estimation. The results consistently demonstrate that parameter recoverability depends strongly on whether the associated biological process is directly observed. Parameters governing microbial dynamics were estimated substantially more accurately when microbial observations were included, whereas parameters associated with oxygen dynamics required dissolved oxygen measurements for reliable estimation. In contrast, interaction parameters linking microbial activity and oxygen dynamics benefited most from combining both observation types. This parameter-specific observation hierarchy remained consistent across calibration experiments, profile likelihood analysis, and Monte Carlo simulations, providing strong evidence that the observed improvements reflect genuine identifiability gains rather than optimization artifacts.

An important methodological finding is that good agreement with decomposition scores alone does not necessarily imply reliable recovery of the underlying biological processes. Although all observation strategies produced similar TADS prediction errors, substantial differences emerged in parameter identifiability, boundary-hit frequencies, and latent-state prediction accuracy. In particular, the profile likelihood and Monte Carlo analyses showed that incorporating complementary biological observations substantially reduced parameter uncertainty while simultaneously improving robustness under measurement noise. These findings highlight the importance of evaluating mechanistic models using identifiability and robustness analyses in addition to conventional goodness-of-fit criteria.

From an operational perspective, the proposed framework aligns naturally with routine forensic practice. Investigators generally have access to decomposition scores, approximate environmental information, and previously documented PMSI estimates, whereas continuous microbiological or biochemical measurements are rarely available. The modular formulation adopted here therefore distinguishes between fixed biological mechanisms, environment-specific parameters, and quantities estimated through calibration. As a result, the same mathematical framework can be recalibrated for different aquatic environments, including coastal waters, lakes, or rivers, without modifying the underlying biological structure.

In the present study, calibration was performed using the published aggregated TADS–PMSI relationship for the Northern Adriatic Sea because individual case-level observations are currently unavailable. Consequently, the estimated parameters should be interpreted as effective descriptors summarizing the combined influence of microbial activity, oxygen dynamics, and environmental conditions within this dataset rather than as unique biological constants. Future availability of longitudinal case-specific measurements will permit more comprehensive identifiability analyses and allow biological variability among different aquatic environments to be quantified explicitly.

Several limitations should also be acknowledged. The current model represents a deliberately reduced description of aquatic decomposition and therefore omits potentially important processes such as scavenger activity, hydrodynamic transport, seasonal ecological variability, and explicit microbial community composition. These simplifications were introduced intentionally to preserve analytical tractability while retaining the dominant mechanisms governing decomposition. The modular structure of the model nevertheless allows such processes to be incorporated naturally as additional state variables or forcing terms when sufficiently detailed datasets become available.

Overall, the proposed framework establishes a transparent and extensible mathematical foundation for mechanistic PMSI estimation. Rather than replacing conventional forensic assessment, it provides a quantitative decision-support tool capable of integrating decomposition observations with environmental information in a biologically interpretable manner. As richer longitudinal datasets become available, the framework offers a systematic basis for improving parameter estimation, uncertainty quantification, and the development of standardized mechanistic approaches for aquatic forensic investigations.

## Declaration of Competing Interest

The authors declare that they have no known competing financial interests or personal relationships that could have appeared to influence the work reported in this paper.

## References

[1] A. A. Vass, The Elusive Universal Post-Mortem Interval Formula, Forensic Science International, 204 1–3 (2011) 34–40.

[2] G. S. Anderson and L. S. Bell, Impact of Marine Submergence and Season on Faunal Colonization and Decomposition of Pig Carcasses in the Salish Sea, PLoS ONE, 11 3 (2016) e0149107.

[3] V. Heaton, A. Lagden, C. Moffatt, T. Simmons, Predicting the Postmortem Submersion Interval for Human Remains Recovered from U.K. Waterways, Journal of Forensic Sciences, 55 2 (2010) 302–307.

[4] C. Palazzo, G. Pelletti, P. Fais, A. Giorgetti, R. Boscolo-Berto, R. M. Gaudio, F. Pirani, A. Tagliabracci, S. Pelotti, Application of Aquatic Decomposition Scores for the Determination of the Post Mortem Submersion Interval on Human Bodies Recovered from the Northern Adriatic Sea, Forensic Science International, 318 (2021) 110599.

[5] M. A. van Daalen, D. S. de Kat, B. F. L. Oude Grotebevelsborg, R. de Leeuwe, J. Warnaar, R. J. Oostra, W. L. J. M. Duijst-Heesters, An Aquatic Decomposition Scoring Method to Potentially Predict the Postmortem Submersion Interval of Bodies Recovered from the North Sea, Journal of Forensic Sciences, 62 2 (2017) 369–373.

[6] C. Palazzo, G. Pelletti, P. Fais, R. Boscolo-Berto, F. Fersini, R. M. Gaudio S. Pelotti, Postmortem Submersion Interval in Human Bodies Recovered from Fresh Water in an Area of Mediterranean Climate: Application and Comparison of Preexisting Models, Forensic Science International, 306 (2020) 110051.

[7] G. Reijnen, H. T. Gelderman, B. F. L. Oude Grotebevelsborg, U. J. L. Reijnders, W. L. J. M. Duijst, The Correlation Between the Aquatic Decomposition Score (ADS) and the Postmortem Submersion Interval Measured in Accumulated Degree Days (ADD) in Bodies Recovered from Fresh Water, Forensic Science, Medicine and Pathology, 14 3 (2018) 301–306.

[8] J. Dalal, S. Sharma, T. Bhardwaj, S. K. Dhattarwal, Assessment of Postmortem Submersion Interval Using Total Aquatic Decomposition Scores of Drowned Human Cadavers, Journal of Forensic Sciences, 68 2 (2023) 549–557.

[9] S. López-Lázaro and C. Castillo-Alonso, Accuracy of Estimating Postmortem Interval Using the Relationship Between Total Body Score and Accumulated Degree-Days: A Systematic Review and Meta-analysis, International Journal of Legal Medicine, 138 (2024) 2659–2670.

[10] M. S. Megyesi, S. P. Nawrocki, N. H. Haskell, Using Accumulated Degree-Days to Estimate the Postmortem Interval from Decomposed Human Remains, Journal of Forensic Sciences, 50 3 (2005) 618–626.

[11] B. M. Dawson, M. Ueland, D. O. Carter, D. McIntyre, P. S. Barton, Bridging the Gap Between Decomposition Theory and Forensic Research on Postmortem Interval, International Journal of Legal Medicine, 138 2 (2024) 509–518.

[12] D. H. Smith, N. Nisbet, C. Ehrett, C. I. Tica, M. M. Atwell, and K. E. Weisensee, Modeling Human Decomposition: A Bayesian Approach, Forensic Science International, 367 (2025) 112309.

[13] C. Bartgis, A. M. LeBrun, R. Ma, L. Zhu, Determination of Time of Death in Forensic Science via a 3-D Whole Body Heat Transfer Model, Journal of Thermal Biology, 62 (2016) 109–115.

[14] D. Justić, N. N. Rabalais, R. E. Turner, W. J. Wiseman Jr., Seasonal Coupling Between Riverborne Nutrients, Net Productivity, and Hypoxia, Marine Pollution Bulletin, 26 4 (1993) 184–189.

[15] J. F. Gillooly, J. H. Brown, G. B. West, V. M. Savage, E. L. Charnov, Effects of Size and Temperature on Metabolic Rate, Science, 293 5538 (2001) 2248–2251.

[16] J. H. Brown, J. F. Gillooly, A. P. Allen, V. M. Savage, G. B. West, Toward a Metabolic Theory of Ecology, Ecology, 85 7 (2004) 1771–1789.

[17] M. T. Madigan, K. S. Bender, D. H. Buckley, W. M. Sattley, D. A. Stahl, Brock Biology of Microorganisms, 15th ed., Pearson, New York, (2018).

[18] J. R. Wallace, J. P. Receveur, P. H. Hutchinson, S. F. Kaszubinski, H. E. Wallace, M. E. Benbow, Microbial community succession on submerged vertebrate carcasses in a tidal river habitat: Implications for aquatic forensic investigations. Journal of Forensic Sciences, 66 (2021) 2307–2318.

[19] C. Jangid, K. Kumari, R. Joshi, M. Hamza, J. Dalal, Estimation of postmortem submersion interval based on microbial community composition in human remains recovered from aquatic environments, Journal of Forensic Sciences, (2026).

[20] M. S. Bone,T. P. R. A. Legrand, M. L. Harvey, M.L. Wos-Oxley, A. P. A. Oxley, Aquatic conditions and bacterial communities as drivers of the decomposition of submerged remains. Forensic Science International, 361 (2024) 112072.

[21] Bray, S. K., Bone, M., Harvey, M. L., Conlan, X. A. Water quality as an indication of postmortem submersion interval (PMSI) for remains in small, isolated fresh water bodies. Australian Journal of Forensic Sciences, 58 (2026) 296–309.

